# A novel genetically distinct *Amdoparvovirus* in *Sorex araneus* in the United Kingdom highlights an unexplored ancestral link

**DOI:** 10.64898/2026.02.04.703478

**Authors:** Tiernan Briggs, Dan Maskell, Dan Henderson, Courtney Graham, Bill Mansfield, David Jorge, Audra-Lynne Schlachter, Matthieu Bernard, Rebecca Callaway, Damian Osmond, Joan Amaya-Cuesta, Florian Pfaff, Henry Ashpitel, Graham Smith, Yogesh Kumar Gupta, Lorraine M McElhinney, Mirjam Schilling

## Abstract

Amdoparvoviruses have historically been documented almost exclusively in carnivores, with recent detections in bats. However, endogenous viral elements in rodent genomes suggest a more ancient and taxonomically broader evolutionary history. Despite this, small mammals have never been systematically surveyed for extant amdoparvovirus infections.

In this study, we used whole genome sequencing to screen four different shrew species and wild American mink in the UK, which may act as a reservoir host for amdoparvoviruses.

We identified a highly divergent *amdoparvovirus* in native common shrews (*Sorex araneus*) from northern England, tentatively named Shrew parvovirus 1(SP 1). Classical *amdoparvovirus* sequences were also detected in wild American mink (*Neogale vison*), confirming the presence of known amdoparvovirus strains in UK mustelids.

Phylogenetic analysis revealed that the shrew virus, SP 1, forms a distinct clade, suggesting ancient divergence or long-term cryptic circulation in small mammal reservoirs. These findings fundamentally challenge the hypothesis that amdoparvoviruses are carnivore-restricted pathogens and underscore the importance of systematic wildlife surveillance for understanding viral host range evolution and assessing spillover risks.

## Introduction

Advances in whole genome sequencing have transformed wildlife surveillance enabling the discovery of novel viruses in unexpected hosts. It is now possible to characterise wildlife viromes in unprecedented detail, offering essential insights into virus diversity and evolution, reservoir hosts, and potential spillover pathways (1).

Amdoparvoviruses (family *Parvoviridae;* subfamily: *Parvovirinae;* genus *Amdoparvovirus*) are single-stranded DNA viruses and have historically been studied almost exclusively in carnivores (2-11). The best-studied member, Aleutian mink disease virus (AMDV), causes chronic immune dysregulation and high mortality in both farmed and wild mink (12). Since its discovery, amdoparvoviruses have been identified in foxes, wildcats, badgers, and other carnivores (2-4, 9, 13), with recent detections also reported in bats (9, 11). This taxonomic distribution has led to the prevailing assumption that amdoparvoviruses are primarily carnivore-adapted pathogens. This however provides a tempting avenue of exploration in which amdoparvoviruses may also be found within small mammals and prey animals, as is suggested by bat and partial rat sequences recently discovered (9, 11, 14).

However, indirect evidence suggests a more ancient and taxonomically broader distribution. The discovery of endogenous viral elements in rodent genomes that are genetically linked to amdoparvoviruses gave rise to speculations about an ancient evolutionary origin and a host range far wider than previously assumed (15). Despite this evolutionary hint, small mammals including shrews have never been systematically surveyed for amdoparvoviruses, limiting our understanding of amdoparvovirus ecology and spillover pathways.

Given their abundance, ecological role, and established capacity as viral reservoirs, shrews represent an ideal but overlooked candidate host for surveying amdoparvovirus diversity beyond carnivores. Shrews have become increasingly recognised as important viral reservoirs of zoonotic and emerging viruses (16-22). This includes the novel Langya virus, a parahenipavirus that has been linked to human disease in both China and Korea (16, 20) and Borna disease virus, which has caused fatal encephalitis in Germany (17). In the UK the relatively recent arrival of the invasive greater white-toothed shrew (*Crocidura russula*) raised ecological and epidemiological concerns (23) as the species’ expansion into Ireland has been associated with the displacement of native shrews and the potential introduction or amplification of pathogens (19). These growing concerns highlight the need for systematic, national wildlife-based surveillance to detect novel or cryptic viral threats (24). To address this knowledge gap, we investigate whether amdoparvoviruses occur beyond their recognized carnivore host range. Utilising metagenomic sequencing, we compared the viromes of four different native and invasive shrew species (*Sorex araneus, S. minutus, Neomys fodiens, C. russula*), from four different regions in England. We additionally sampled sympatric American mink (*Neogale vison*), an established amdoparvovirus host, to contextualize any shrew-associated findings and assess potential spillover risks.

We discovered and characterised a previously undescribed amdoparvovirus in *S. araneus* from northern England. Our analyses reveal the presence of a highly divergent amdoparvovirus lineage in UK shrews and provide the first evidence of such a virus in this host group. These findings challenge previous research showing amdoparvoviruses are primarily restricted to carnivores. This amdoparvovirus could further represent a lineage that may have long circulated undetected in small mammalian reservoirs and might provide a crucial link in understanding the evolutionary history of amdoparvoviruses.

## Results

### A novel *amdoparvovirus* discovered in Eurasian shrews *(S. araneus)*

To investigate the presence of amdoparvovirus in UK shrew populations, metagenomic sequencing of samples from both native and invasive shrews collected over two years were analysed. The samples represent three sites in England, including two locations where greater white toothed shrews (*C. russula*) are established **(Figure 1. A)** (23). Data was generated using short-read sequencing of 47 animals (*S. araneus* n = 11, *S. minutus* n = 3, *N. fodiens* n = 1, *C. russula* n = 32), yielding 376 tissue samples. Shrew parvovirus 1 was detected by both sequencing and PCR in 5/11 *S. araneus* individuals **(Figure 1. B)** across multiple organs suggesting a systemic infection within these individuals. SP 1 was notably absent within all other species of shrews tested and was only found within *S. araneus* animals from County Durham **(Figure 1.B & C)** over both years where shrews were collected (n=3 for year one and n=2 for year two).

**Figure 1:**
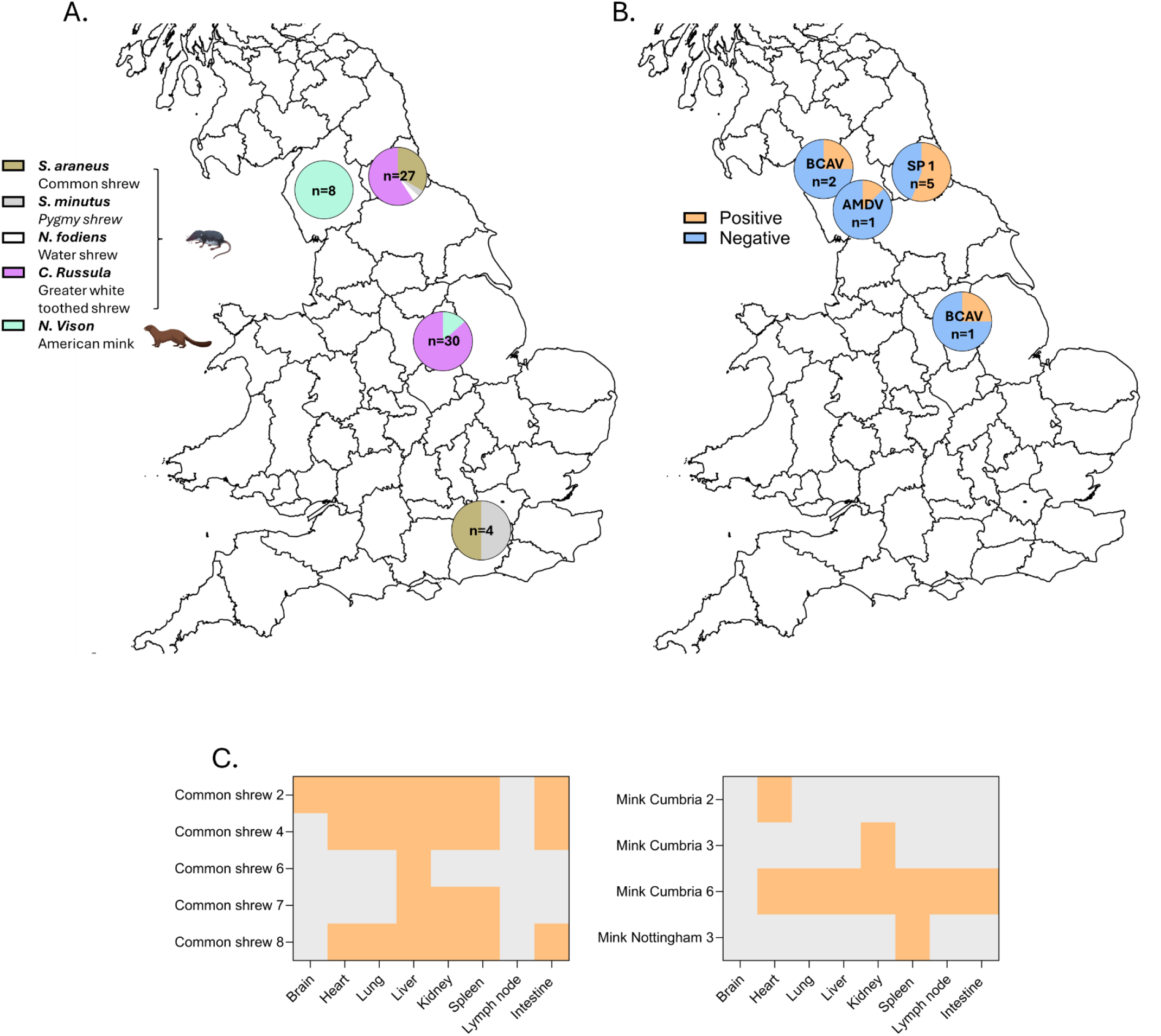
Sampling areas show spatial distribution of amdoviruses within the UK as well as within different tissues. **A. Animals were sampled in four different areas in England**, where possible these overlapped. **B. Amdovirus distribution was greater in shrews than mink.** SP 1 was detected in over half of the common shrews whilst the mink did not show this distribution. **C. SA-LV 1 detection appears systemic within shrews whilst classical amdoviral sequence detection is more limited in mink**, in contrast to all but one mink. **Abbreviations: SP 1;** Shrew parvovirus 1, **AMDV;** Aleutian mink disease virus, **BCAV;** British Columbia amdovirus. Created in BioRender. https://BioRender.com/wwyrc2w.

Mink samples (*N. vison*, n=12, 84 tissues) were also screened to determine the prevalence of amdoparvovirus within UK mustelid populations **(Figure 1. A)**. Metagenomic sequencing identified two distinct amdoparvovirus sequences **(Figure 1. B)**. Unlike the newly identified amdoparvovirus in shrews, the mink-associated sequences were restricted to fewer tissues, primarily the spleen, with the exception of one individual where AMDV was detected systemically **(Figure 1. C)**.

### Phylogenetic characterisation of SP 1 sequence reveals a distinct amdo-like lineage

To determine the evolutionary placement of the detected novel shrew amdoparvovirus, we compared its genome to known amdoparvoviruses. This revealed a nucleotide sequence similarity of 70% to AMDV, with its closest known relative being Bat parvovirus 11 with a nucleotide similarity of 70% to SP 1 (11) **(Figure 2. B)**. Following ICTV guidelines *(25)*, the sequence was designated Shrew parvovirus 1 (SP 1) due to a 52% amino acid NS1 protein sequence identity compared to AMDV. Entire genomes were retrieved from 4 animals, the remaining being fragmented with an average coverage of 1000 reads per region. A consensus sequence was used for the following analyses.

**Figure 2:**
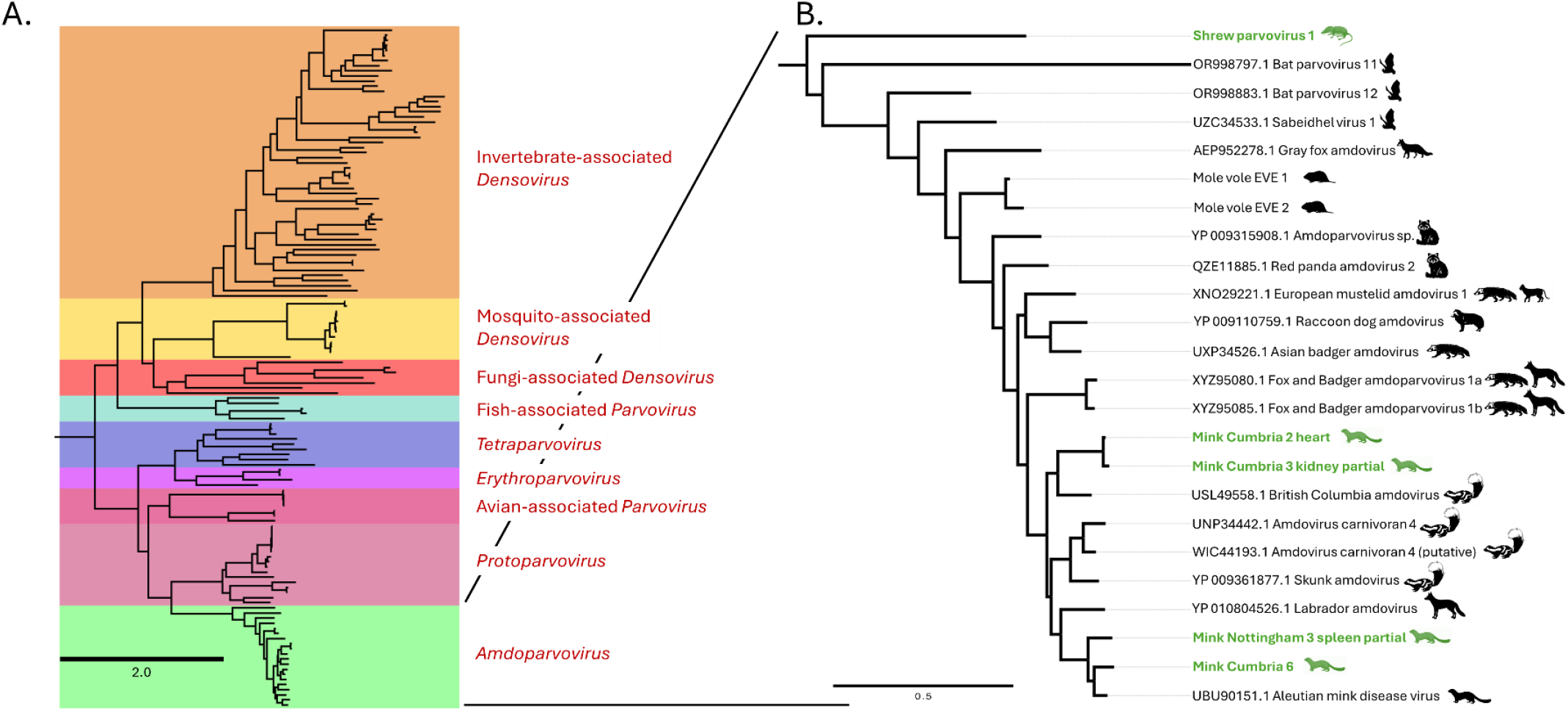
Shrew amdo-like virus 1 shows a large divergence from classical amdoviruses which are also detected within British mustelids. **A. Whole family parvoviridae tree** rooted to Bos taurus papillomavirus 1 E1 protein (NP 056739). 1000 bootstraps, Model: Q.pfam+F+I+R7. **B. Amdoparvovirus tree** generated from only amdoparvovirus sequences. Rooted to Canine parvovirus NS1 protein (WQF67428.1). SP 1 sequences collapsed into single node for simplicity. Ancient mole-vole sequences derived from mole-vole genome outlined in Penzes et al 2018. 1000 bootstraps, Model: Q.INSECT+F+G4. **Abbreviations; EVE;** endogenized viral element. Sequences used were Ref-Seq, available from NCBI, last accessed 22/08/2025. Created in BioRender https://BioRender.com/xgnenkf.

Phylogenetic analysis placed SP 1 in a distinct clade within the *amdoparvovirus* genus **(Figure 2. A)**. The NS1 **(Figure 2. B)** gene sequence tree placed SP 1 on an independent branch, separate from mustelid amdoparvoviruses, but more closely aligned with bat-associated amdoparvoviruses and other non-mustelid sequences. Conversely, VP1 sequence analysis grouped SP 1 with sequences of labrador amdoparvovirus 2 and a partial sequence of rat parvovirus **(Supp. 1 & 2)**. However, due to a higher level of VP1 amino acid sequence conservation of 69% compared to AMDV, taxonomic demarcation relies on the NS1 gene sequence (25).

In contrast, the amdoparvovirus sequences obtained from mink clustered within the known mustelid amdoparvovirus group, showing close relationships with AMDV and British Columbia amdoparvovirus (BCAV). The BCAV-like sequences displayed greater divergence, suggesting a possible UK-specific lineage given their distribution across multiple counties **(Figure 2. B)**.

Alongside the sequences acquired from UK animals, a data mining approached was attempted in order to expand our understanding of the geographical range of SP 1. However, no closely related sequences could be retrieved, highlighting the gap in published amdoparvoviral genomes. Partial sequences from rodents and carnivore species were detected in line with previously reported viral genomes **(Supp. 2 A & B)**. Overall, these findings in shrews expand the known host range of amdoparvoviruses beyond carnivores and bats. Our analysis also highlights the phylogenetic proximity of SP 1 to both bat amdoparvoviruses and ancient parvoviral elements in the Transcaucasian mole vole (*Ellobius lutescens*) genome derived from Penzes, Marsile-Medun (15). This suggests that amdoparvoviruses may have an ancient evolutionary origin predating the diversification of modern mammalian hosts.

### Validation of SP 1 using qPCR screen

To enable reliable and scalable screening for SP 1 in future surveillance efforts, we designed and validated primers targeting the NS1 region based on the *de novo* assembled sequence. PCR positive samples mirrored the SP 1 distribution pattern observed in our metagenomic data validating the qPCR we have established. Amplicons were verified by Sanger sequencing, which matched the assembled SP 1 genome, confirming the presence of a highly divergent *amdoparvovirus* across multiple organs in shrews, consistent with systemic infections.

### Characteristic amdoparvoviral protein motifs are retained despite low nucleotide sequence homology

Comparative motif analysis of the SP 1 genome revealed the preservation of hallmark amdoparvoviral domains, including rolling circle replication motifs II and III and the SF3 helicase region. Despite low overall nucleotide identity, these motifs are conserved on amino acid level, with the rolling circle replication II motif showing homology to other amdoparvoviruses **(Figure 3. A)**.

**Figure 3:**
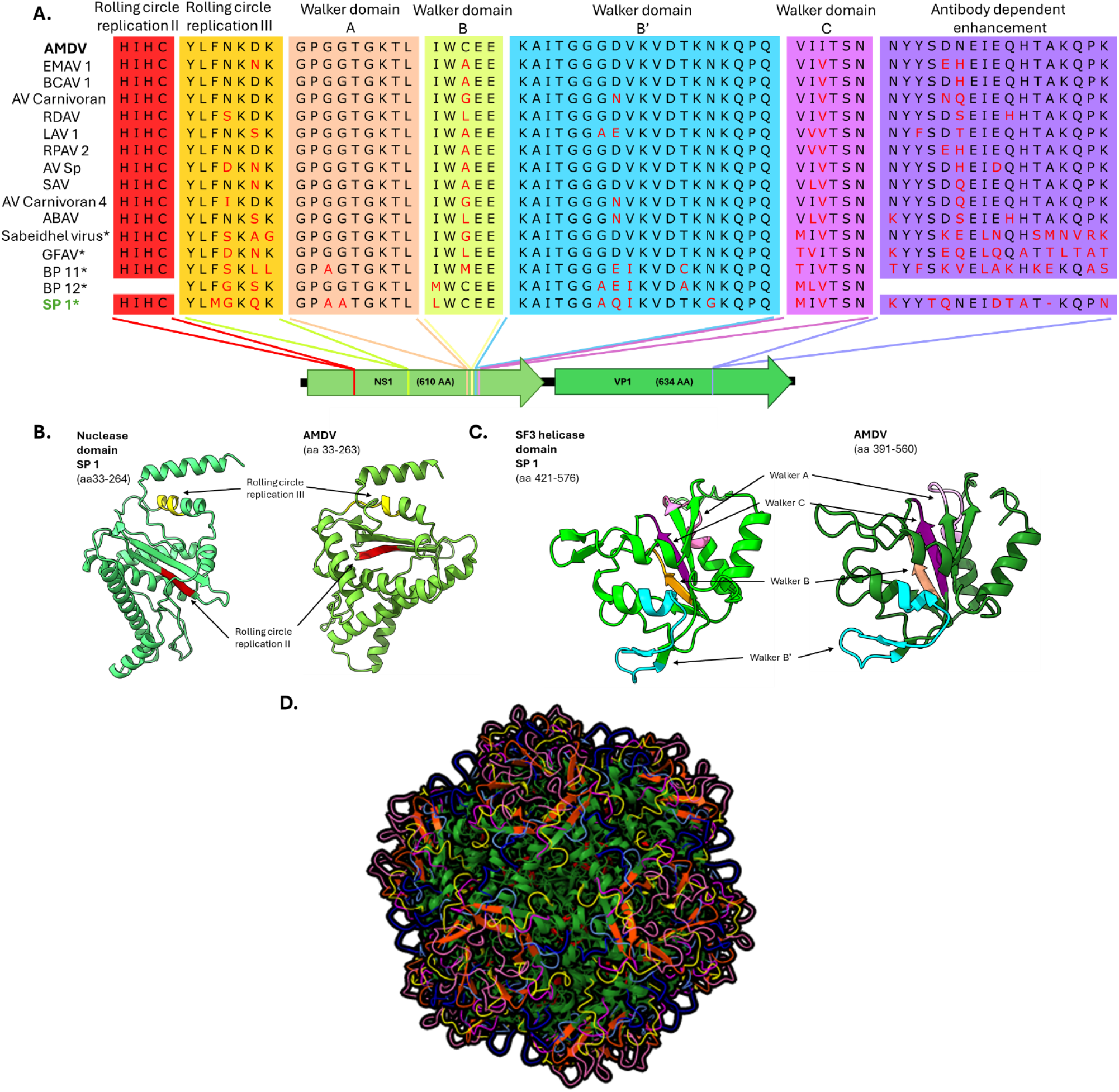
Putative shrew amdo-like virus 1 motifs are structurally conserved despite little amino acid sequence conservation. **A. Motif conservation**, common ADV protein motifs were aligned with SA-LV1. Species with stars denote non-mustelid hosts. **B & C. Protein structure predictions** Key protein domains are similar to previously known structures; domains are coloured with respect to above figure **(A). D. Full capsid modelling indicates SA-LV1 icosahedral structure**, surface loops were coloured based upon visual cues. **Abbreviations; AMDV**: Aleutian mink disease virus, **EMAV**: European mustelid amdoparvovirus, **BCAV1**: British Columbia amdovirus 1, **AV carnivoran (4)**; Amdovirus carnivoran (4), **RDAV**; Raccoon dog amdovirus, **GFAV**; Gray fox amdovirus, **AV Sp**; Amdovirus species, **SAV**; Skunk amdovirus, **ABAV**; Asian badger amdovirus, **LAV**; Labrador amdovirus, **BP (11/12)**; Bat parvovirus (11/12), **SP 1;** Shrew parvovirus 1, **SF3; Super family 3.** Sequences used were Ref-Seq, available from NCBI, last accessed 22/08/2025.

The VP1 protein contains the antibody-dependent enhancement (ADE) region described in other amdoparvoviruses (5), although with low amino acid sequence identity as also shown for other amdoparvovirus species present outside of mustelid and carnivore hosts such as the gray fox and sabeidhel amdoparvoviruses. The NS1 polyprotein contains an HUH superfamily endonuclease and a superfamily 3 helicase. AlphaFold based structural prediction (26), supports nuclease and helicase domain topology conservation, indicating that sequence divergence had not disrupted key replication associated structures seen within all amdoparvovirus species **(Figures 3. B&C)**. Capsid modelling predicted a typical icosahedral morphology **(Figure 3. D)** consistent with other Amdoparvovirus species. However, the VP1 surface loop regions were notably divergent, indicating structural variation. These findings confirm that SP 1 retains key amdoparvovirus features despite high sequence divergence, supporting its classification and expanding the known host range beyond carnivores and bats.

### Histological examinations confirm systematic infection but no overt pathology

Histological examination of tissues revealed mild multifocal inflammation characterised by peribronchiolar and perivascular lymphohistiocytic infiltrates in the lungs, and minimal multifocal periportal lymphohistiocytic infiltrates in the liver, in two of the three positive shrews examined **(Figure 4. A-D)**. Concurrently, in one of these two shrews, minimal multifocal granulocytic infiltrates were infrequently observed in the intestinal submucosa, with evidence of intraluminal nematode larva, and tentative coccidian merozoites, consistent with parasitic inflammation. However, no lesions suggestive of viral infection were detected.

**Figure 4:**
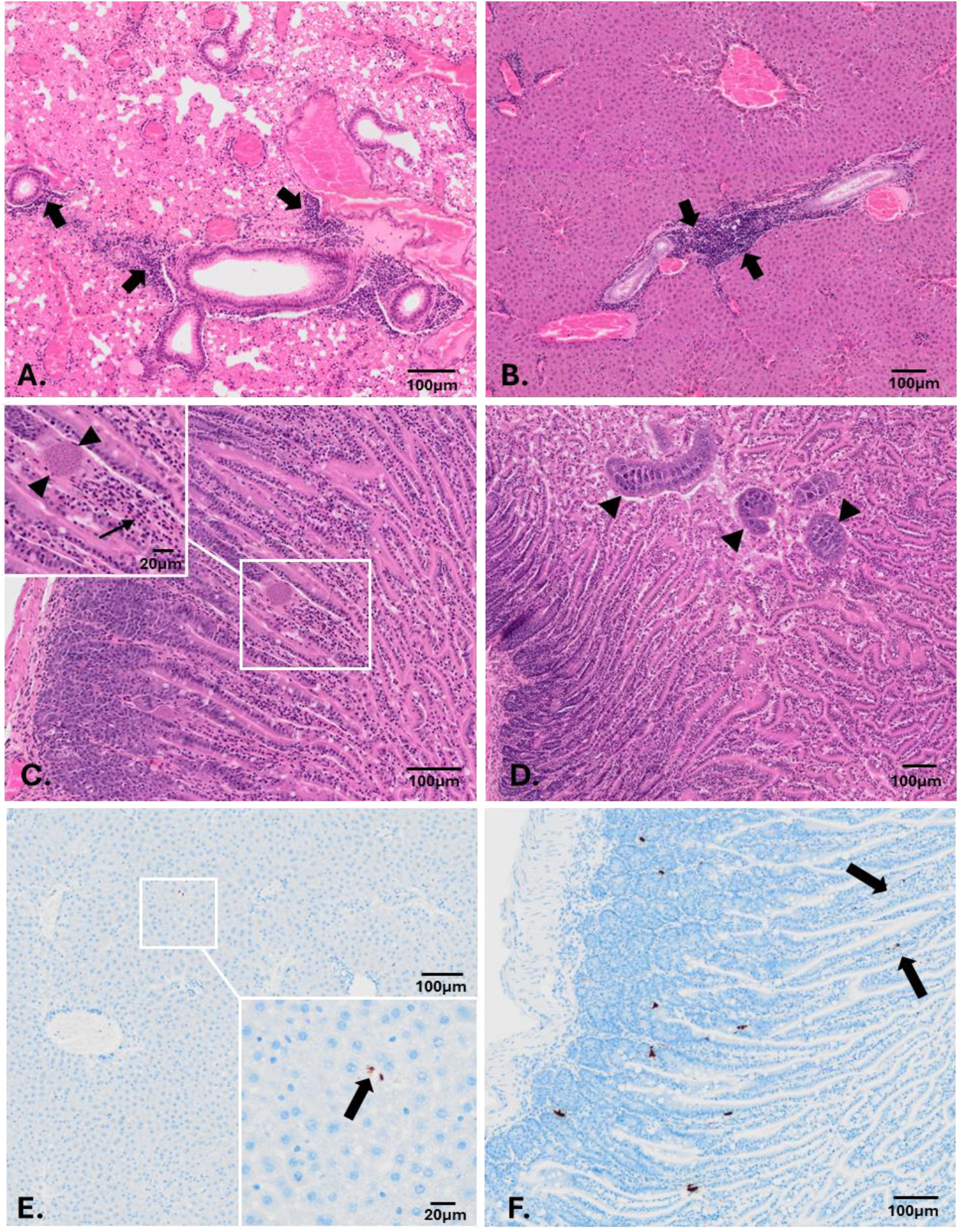
SA-LV 1 does not cause overt pathology in Sorex Araneus. Representative H&E sections of A. lung, B. liver, and C & D. small intestine, from one S. araneus. No histopathological lesions attributable to viral infection were observed in all sections examined. Mild multifocal lymphohistiocytic peribronchiolar and periportal infiltrates were present in the lung and liver respectively (short arrows). Mild multifocal granulocytic infiltrates (thin arrow) were seen alongside c) coccidian (arrowheads) and d) nematode (arrowheads) infestation, in intestinal sections. **cISH labelling revealed SA-LV1 nucleic acid within E. liver and F intestinal tissues**. Scale bars are shown. **Abbreviations: cISH;** Chromogenic in situ hybridisation. **H & E;** Haematoxylin & Eosin. **SA-LV 1;** Shrew amdovirus-like 1

Chromogenic in situ hybridisation (cISH) detected SP 1 DNA in the liver and intestine of the same individual **(Figure 4. E & F)** displaying histological changes, while all other samples examined from virus positive shrews were negative. Negative DapB controls **(Suppl. 3)** provided evidence of the specificity of staining, location of labelling was not associated with any cell type and often found extracellularly.

## Discussion

It has long been speculated that members of the genus *Amdoparvovirus* may infect hosts beyond their typical mustelid and carnivore range (5). Here, we report for the first time the detection of an amdoparvovirus in UK common shrews (*S. araneus*), tentatively named shrew parvovirus 1 (SA-LV1). Our findings confirm the hypothesis that these viruses are not restricted to their usual mustelid and carnivore hosts (5), as suggested by the recent detections of bat-borne amdoparvoviruses (9, 11).

This discovery provides a crucial link in understanding the broader host spectrum and evolutionary history of *amdoparvovirus*, revealing a potential lineage that may have long circulated undetected in small mammalian reservoirs.

Genomic classification and characterisation indicate that SP 1 falls within the *amdoparvovirus* genus but is highly divergent, sharing 70% nucleotide sequence identity and 10% coverage with its nearest known relative, bat parvovirus 11, which is itself a highly diverged species (11). The extent of sequence divergence suggests that SP 1 represent a broader viral lineage that may have originated in prey species. According to ICTV guidelines, amdoparvoviruses exhibit species demarcation thresholds of approximately 85% sequence identity. Therefore, SP 1, with a 52% NS1 sequence homology to AMDV represents a novel and distinct lineage. Phylogenetic analyses of the NS1 and more conserved VP1 proteins place SP 1 within the *amdoparvovirus* clade (25, 27), though low bootstrap support and topological instability likely reflect its evolutionary distance from other known members. The retention of conserved nuclease, SF3 helicase, and other hallmark amdoparvoviral genomic features (25, 27) supports its inclusion within the genus.

The detection of SP 1 was limited to a single UK region, near Durham, and its broader distribution remains unknown. Whilst we also identified the classical *amdoparvovirus* species AMDV and BCAV in mustelids in the UK, no mink samples from Durham were available for this study. Consequently, the prevalence of SP 1 in these animals is unknown. Given the protected status of *S. araneus*, expanded surveillance depends on citizen science initiatives and opportunistic submissions of shrews found dead, or studies conducted under Natural England licences, limiting our understanding of viruses circulating in this species. Although we were able to test native and invasive shrews from different areas within this study, we did not detect any *amdoparvovirus* sequences outside the Durham region, possibly explained due to smaller sample sizes. Interestingly, no detection of SP 1 was made in the greater white-toothed shrews from he same area, suggesting hitherto limited spillover of microbes between the native and invasive species. Nonetheless, the degree of sequence divergence from other amdoparvoviruses suggests that related viruses may be circulating more widely among native UK shrews and other small mammal populations.

Despite extensive nucleic acid and amino acid sequence divergence, structural and motif conservation indicates retention of protein structural similarities. Alphafold prediction revealed well-preserved domains and topology similar to previously reported members of the species (5). Though notable shifts in surface loop positioning and the ADE motif, implicated in host cell attachment (28) suggest host-specific adaptations, a pattern mirrored in other non-carnivore amdoparvoviruses such as bat parvovirus 11 and sabeidhel virus (9, 11). This supports the hypothesis that functional constraints shape amdoparvovirus evolution across diverse hosts.

Histopathological examination of infected shrews revealed no lesions attributable to viral infection. Whilst mild pulmonary inflammation and hepatic changes were detected, these were non-specific and coincided with intestinal parasitism. In addition, changes were not colocalised with cISH labelling and cannot confidently be linked to viral presence. Although positive labelling was detected by cISH in one positive shrew, the specific cell type involved could not be identified and labelling was often extracellular. Furthermore, the lack of positive control material makes the significance of positive labelling unclear. It is also possible that due to the practicalities of field investigation, as tissue samples were processed up to 48 hours postmortem, autolysis could mask subtle pathology, and result in non-specific or extracellular labelling. Therefore, while changes may represent a non-pathogenic SP 1 or persistent systemic infection in shrews, further investigation with fresh tissue, positive control material and a greater sample size is needed.

The absence of histopathologic findings contrasts with AMDV in mink which, dependent on virus strain, dose and age at the time of infection, can result in multiple clinical syndromes. In kits, acute fulminant interstitial pneumonia with varying morbidity and mortality is seen, whereas in adults, persistent chronic infection leads to immune dysregulation and ultimately immune complex disease, resulting in arteritis and glomerulonephritis (6-8, 12, 29-32).

The evolutionary origin of SP 1 remains unresolved as data-mining of global metagenomic datasets using SRAminer (33) did not reveal related sequences or fragments. However, the increasing amount of publicly available SRA datasets together with some more targeted surveillance projects in the future will help to elucidate the evolutionary past of amdoparvoviruses. Our discovery challenges the long-held hypothesis that amdoparvoviruses evolved solely within carnivores raising the possibility that prey species such as shrews or rodents may represent ancestral hosts. Alternatively, this divergent lineage could have emerged through historic host exchange during the fur trade era in Britain. Future work should expand surveillance across small mammal populations.

## Materials & methods

### Sample collection

Trap sites / areas for shrews were chosen based on proximity to original detections of non-native *C. russula* (23), suitability of habitat and accessibility at short notice. Between 4 and 60 Longworth traps, baited with wild bird seed and casters, were set at each trap site. Traps were in place for sessions of 1-4 trapping nights and checked at least every 12 hours.

Shrews were despatched by cervical dislocation. All operatives were appropriately trained for trapping shrews to comply with the Natural England general licence.

Shrews sampled in the Surrey area were kindly provided by local pet cats and citizen science efforts.

Cage traps for mink were fitted with a remote monitoring device and mink scent-soaked filter balls placed behind the footplate. Traps were then set up on floating rafts and deployed on the waterway. Mink were humanely despatched in line with UK legislation and the Wildlife and Countryside Act 1981 and categorised based on sex, size, weight, length and coat colour. Tissue samples were taken for genetic analysis.

### Nucleic acid extraction

Shrew carcasses were dissected, and the brain, lung, heart, liver, kidney, spleen, intestine and presumptive cervical lymph nodes were removed with half of the sample being collected for histopathlogy and the other half stored at -80 °C. The stored samples were used for total nucleic acid extraction; individual tissues were homogenised in 600 µl 1M PBS using a 5mm steel bead and the (Invitrogen bead mill). A Kingfisher ^™^extraction was performed on 200 µl of the crude homogenate and extracted into 90 µl of extraction buffer in line with the Kingfisher ^™^ MagMax protocol (Thermofisher).

### cDNA synthesis

cDNA synthesis was performed using 1^st^ strand Superscript IV (Thermofisher) and 2^nd^ strand (NEB). A master mix including 1 µl random hexamers was prepared with 8.5 µl of RNA/DNA added to 4.5 µl of master mix according to manufacturer’s protocol. The plate was transferred to a thermal cycler at 65 °C for 5 minutes to allow primer annealing. The RT master mix was prepared in line with manufacturer’s instructions and added to the sample plate. First strand synthesis was performed at 23 °C for 10 minutes, 55 °C for 10 minutes and finally 80 °C for 10 minutes. Second strand synthesis was performed using NEBNext Ultra II with master mixes composed according to manufacturer’s instructions. 60 µl of master mix was added to each sample and cooled to 16 °C for 60 minutes. Samples were purified using Agencourt Ampure XP in line with the manufacturer’s instructions and eluted in 10 µl 10 mM Tris (pH 7.5).

### Whole genome sequencing

DNA concentrations were read using the Fluostar Omega Plate reader (BMG Labtech) using QuantiFluor reagents (Promega) in line with the manufacturer’s protocols for sample and standard preparation and sample concentrations were normalised to between 0.2 and 1 ng/µl.

Sequencing libraries were prepared using the Nextera XT kit (Illumina, Cambridge, UK) and analysed on a NextSeq sequencer (Illumina, Cambridge, UK) with 2 × 150 base paired end reads.

### Bioinformatic assemblies

Following on from Illumina sequencing raw fastq files were quality controlled using FastP (34) with automatic adapter detection, minimum length filtering of 31 and a phred-cutoff of 4 and taxonomically classified with Kraken 2 (35) and passed through to Krakentools (36) to facilitate the removal of bacterial and bacteriophage reads. Host reads were removed with BWA-MEM by alignment to genomes (*S. araneus* GCF_027595985.1 and *N. vison* GCF_020171115.1), unaligned reads were extracted with SAMtools (37), non-host reads were assembled using metaspades (38) with kmer lengths of 21,41,71,101,127 and scaffolded with Lilypad (Part of the BBMap suite). Putative viral contigs were identified through ViralVerify (34, 39, 40) and analysed by BLAST using a custom blastn (41) database of all NCBI ref-seq viruses and Pfam-A (42). Quality control was performed with QUAST (43) pre and post scaffolding.

Amdparvovirus related EVE’s were identified using TBLASTN searches of the Transcaucasian Mole Vole (*E. lutescens*) genome (GCA_001685075.1) using amdoparvovirus sequences (UBU90151.1 & AEK27534.1) with matching genomic regions extracted as nucleotide sequences using BEDtools (44).

### Phylogenetic analysis

Extracted amdoparvovirus and parvovirus contigs were analysed through ORFfinder to generate NS1 protein sequences and then aligned using MAFFT (45) default settings. IQTREE2 (46) was used to generate a maximum likelihood tree of both parvovirus and amdoparvoviral sequences using automatic model selection and 1000 ultrafast bootstraps and visualised with FigTree.

### Motif analysis

NS1 sequences for SP 1 and previously reported reference amdoparvovirus sequences were aligned with MAFFT (45) and common amdoparvovirus protein motifs were highlighted.

### Protein structure analysis

NS1 protein domains were generated using InterPro (47) to extract the nuclease and helicase domains of SP 1 and AMDV structural analysis was performed using ColabFold (26) and annotations made with ChimeraX (48). VP1 protein structures were estimated using AlphaFold3 (49) and a whole capsid model was generated using ChimeraX (48).

### SRAminer

For the SRAmining approach we used the pipeline available from https://gitlab.com/FPfaff/sraminer. The workflow includes SRA database download,

Blastx of database with sequence of interest and reporting on matches to this. A description of the complete method is available in Eshak et al. (33).

### Histopathology

Tissue samples were fixed in buffered formalin, blocked and processed into paraffin blocks before sectioning. 4 µm sections were taken using a microtome and stained with H & E for examination on a Gemini Autostainer (Epredia) using the following programme.

**Table 1:** H & E Staining protocol. After fixation and embedding sectioned samples were stained using the following automated protocol with a Gemini Autostainer (Epredia). Stained samples were then imaged and processed using FiJI.

| Position | Solution | Time (mm:ss) |
| --- | --- | --- |
| 1 | Xylene | 5 minutes |
| 2 | Absolute Alcohol | 3 minutes |
| 3 | Haematoxylin | 8 minutes |
| 4 | Tap Water | 30 seconds |
| 5 | 1% Acid Alcohol | 20 seconds |
| 6 | Tap Water | 30 seconds |
| 7 | 1.5% Lithium Carbonate | 30 seconds |
| 8 | Tap Water | 30 seconds |
| 9 | 1% Eosin | 50 seconds |
| 10 | Tap Water | 30 seconds |
| 11 | Absolute Alcohol | 30 seconds |
| 12 | Absolute Alcohol | 1 minute |
| 13 | Xylene | 5 minutes |

### cISH

All in situ hybridisation staining was conducted on the Bond RX autostainer (Leica Biosystems) using probe sequences generated from the full-length SP 1 genome (PX921647). Tissue sections were stained following the manufacturer’s instructions. Sections were baked then dewaxed with Bond dewax solution (AR9222, Leica Biosystems) before heat-induced epitope retrieval using Bond Epitope Retrieval Solution 2 (AR9640, Leica Biosystems). Slides were treated with Protease Plus (ACD) and hybridised with the V-APV-NS1-VP1-C1 LS probe (1838378-C1,ACD). Staining was amplified using the LS detection kit (322100, ACD). Labelling was demonstrated using the Bond Polymer Refine Detection kit (DS9800, Leica Biosystems); The post primary was supplemented with normal ovine serum (1/33 (v/v) dilution; Sigma-Aldrich) and normal swine serum (1/33 (v/v) dilution; Vector Laboratories), labelling was visualised using DAB chromogen and counterstained with Mayer’s haematoxylin. Bond Wash Solution (AR9590, Leica Biosystems) was used for rinsing sections between incubations. Sections were then dehydrated, cleared and glass coverslip mounted using ClearVue mountant XYL (Epredia). Sections were visualised by two experienced pathologists using the Olympus VS2000, and negative DapB controls examined to determine the specificity of staining.

### PCR

PCR reactions were set up as follows:

**Table 2:** qPCR reaction reagent conditions. Primers obtained from Sigma (Merck) RT-PCR kit used was AgPath-ID™ One-Step RT-PCR Kit (ThermoFisher).

| Name | 1x (μl) |
| --- | --- |
| DNA | 5 |
| 25x RT-PCR enzyme mix | 1 |
| 2X reaction buffer | 12.5 |
| Forward primer (5 μM) | 2.5 |
| Reverse (5 μM) | 2.5 |
| dH<sub>2</sub>O | 1.5 |

**Table 3:** qPCR reaction running conditions. The PCR reaction was set up as follows, an Aria MX thermocycler was used and products run on a 1% agarose gel to confirm probe detection. Data was collected in real time

| Step | Temperature (°C) | Time | Cycles |
| --- | --- | --- | --- |
| Reverse transcription | 45 | 10 minutes |  |
| Initial denaturation | 95 | 10 minutes |  |
| Denaturation | 95 | 15 seconds | x42 |
| Annealing | 57 (ramp up 0.2/cycle) | 20 seconds |  |
| Elongation | 72 | 30 seconds |  |

### Primer and Probe sequences

**Table 4:**
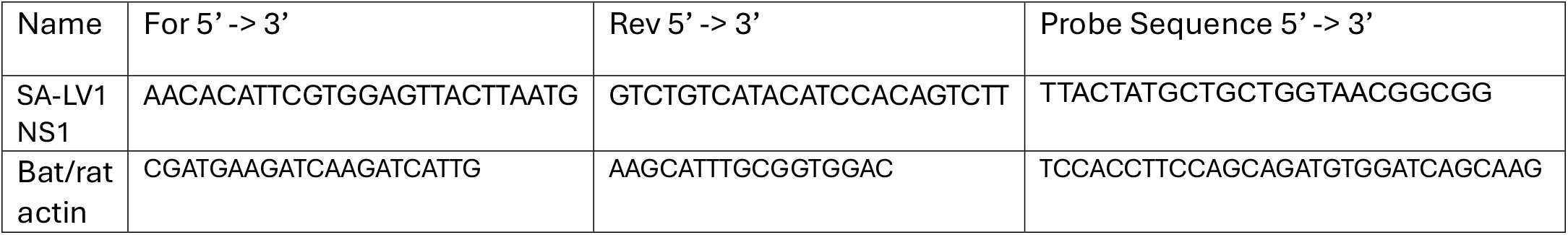
Primer sequences. All primers supplied by Merck dry, eluted in TrisEDTA to a stock concentration of 100mM. Bat/rat actin control primers derived from (50)

## Author contributions

Manuscript first draft: TB; Sampling: DH, CG, BM; Sample preparation: TB, DM, DO, JAC Bioinformatics & molecular characterisation: TB, DM, FP; Pathology & Imaging: DJ, ALS, MB, RC, HA; Project management/supervision; GS, YKG, LMM, MS; Manuscript review & edit: all co-authors.

## Acknowledgments

We would like to thank Cleo & Ptolly who’s hunting expertise helped us greatly in expanding our sample size and allowed us to collect data within Surrey.

We also want to thank the Waterlife Recovery Trust, partners, and funders for their work in collecting mink samples for us to test.

We would also like to thank Matthew Arnold (RVC) for the helpful discussions.

## Data availability

All assembled sequences uploaded to NCBI: Shrew parvovirus 1 - PX921647, Mink sequences – PX930814-PX930820.

SRA data available: PRJNA1418128.

## Ethics statement

The authors confirm that all ethical policies of the journal, noted on the author

## References

1. Shi Y, Li Y, Li H, Haerheng A, Marcelino VR, Lu M, et al. Extensive cross-species transmission of pathogens and antibiotic resistance genes in mammals neglected by public health surveillance. Cell. 2025;188(23):6591–605 e14.

2. Bloom ME, Race RE, Wolfinbarger JB. Characterization of Aleutian disease virus as a parvovirus. J Virol. 1980;35(3):836–43.

3. Canuti M, McDonald E, Graham SM, Rodrigues B, Bouchard E, Neville R, et al. Multi-host dispersal of known and novel carnivore amdoparvoviruses. Virus Evol. 2020;6(2):veaa072.

4. Canuti M, O’Leary KE, Hunter BD, Spearman G, Ojkic D, Whitney HG, et al. Driving forces behind the evolution of the Aleutian mink disease parvovirus in the context of intensive farming. Virus Evol. 2016;2(1):vew004.

5. Canuti M, Penzes JJ, Lang AS. A new perspective on the evolution and diversity of the genus Amdoparvovirus (family Parvoviridae) through genetic characterization, structural homology modeling, and phylogenetics. Virus Evol. 2022;8(1):veac056.

6. Eklund CM, Hadlow WJ, Kennedy RC, Boyle CC, Jackson TA. Aleutian disease of mink: properties of the etiologic agent and the host responses. J Infect Dis. 1968;118(5):510–26.

7. Hadlow WJ, Race RE, Kennedy RC. Comparative pathogenicity of four strains of Aleutian disease virus for pastel and sapphire mink. Infect Immun. 1983;41(3):1016–23.

8. Hadlow WJ, Race RE, Kennedy RC. Temporal replication of the Pullman strain of Aleutian disease virus in royal pastel mink. J Virol. 1985;55(3):853–6.

9. Kamani J, Gonzalez-Miguel J, Msheliza EG, Goldberg TL. Straw-Colored Fruit Bats (Eidolon helvum) and Their Bat Flies (Cyclopodia greefi) in Nigeria Host Viruses with Multifarious Modes of Transmission. Vector Borne Zoonotic Dis. 2022;22(11):545–52.

10. Li L, Pesavento PA, Woods L, Clifford DL, Luff J, Wang C, et al. Novel amdovirus in gray foxes. Emerg Infect Dis. 2011;17(10):1876–8.

11. Wang Y, Xu P, Han Y, Zhao W, Zhao L, Li R, et al. Unveiling bat-borne viruses: a comprehensive classification and analysis of virome evolution. Microbiome. 2024;12(1):235.

12. Vahedi SM, Salek Ardestani S, Banabazi MH, Clark F. Epidemiology, pathogenesis, and diagnosis of Aleutian disease caused by Aleutian mink disease virus: A literature review with a perspective of genomic breeding for disease control in American mink (Neogale vison). Virus Res. 2023;336:199208.

13. Canuti M, Mira F, Villanua D, Rodriguez-Pastor R, Guercio A, Urra F, et al. Molecular ecology of novel amdoparvoviruses and old protoparvoviruses in Spanish wild carnivorans. Infect Genet Evol. 2025;128:105714.

14. Wu Z, Lu L, Du J, Yang L, Ren X, Liu B, et al. Comparative analysis of rodent and small mammal viromes to better understand the wildlife origin of emerging infectious diseases. Microbiome. 2018;6(1):178.

15. Penzes JJ, Marsile-Medun S, Agbandje-McKenna M, Gifford RJ. Endogenous amdoparvovirus-related elements reveal insights into the biology and evolution of vertebrate parvoviruses. Virus Evol. 2018;4(2):vey026.

16. Chakraborty S, Chandran D, Mohapatra RK, Islam MA, Alagawany M, Bhattacharya M, et al. Langya virus, a newly identified Henipavirus in China - Zoonotic pathogen causing febrile illness in humans, and its health concerns: Current knowledge and counteracting strategies - Correspondence. Int J Surg. 2022;105:106882.

17. Ebinger A, Santos PD, Pfaff F, Durrwald R, Kolodziejek J, Schlottau K, et al. Lethal Borna disease virus 1 infections of humans and animals - in-depth molecular epidemiology and phylogeography. Nat Commun. 2024;15(1):7908.

18. Gong HY, Chen RX, Tan SM, Wang X, Chen JM, Zhang YL, et al. Viruses Identified in Shrews (Soricidae) and Their Biomedical Significance. Viruses. 2024:16(9).

19. Nally JE, Arent Z, Bayles DO, Hornsby RL, Gilmore C, Regan S, et al. Emerging Infectious Disease Implications of Invasive Mammalian Species: The Greater White-Toothed Shrew (Crocidura russula) Is Associated With a Novel Serovar of Pathogenic Leptospira in Ireland. PLoS Negl Trop Dis. 2016;10(12):e0005174.

20. Natasha A, Pye SE, Park K, Rajoriya S, Yang I, Park J, et al. Detection and characterization of Langya virus in Crocidura lasiura (the Ussuri white-toothed shrew), Republic of Korea. One Health. 2025;20:101017.

21. Parry RH, Yamada KYH, Hood WR, Zhao Y, Lu JY, Seluanov A, et al. Henipavirus in Northern Short-Tailed Shrew, Alabama, USA. Emerg Infect Dis. 2025;31(2):392–4.

22. Rasche A, Lehmann F, Konig A, Goldmann N, Corman VM, Moreira-Soto A, et al. Highly diversified shrew hepatitis B viruses corroborate ancient origins and divergent infection patterns of mammalian hepadnaviruses. Proc Natl Acad Sci U S A. 2019;116(34):17007–12.

23. Smith GC, Bond IF, Coult T, Henderson D, Graham C, Brand E, et al. Establishment, current status and possible origin of the greater white-toothed shrew Crocidura russula in Great Britain. Biological Invasions. 2025;27(9):197.

24. Gupta YK, Adams I, van Aerle R, Avant J, Bass D, Batista FM, et al. An integrated One Health initiative for pathogen genomic surveillance in the UK. Microb Genom. 2025:11(11).

25. Cotmore SF, Agbandje-McKenna M, Canuti M, Chiorini JA, Eis-Hubinger AM, Hughes J, et al. ICTV Virus Taxonomy Profile: Parvoviridae. J Gen Virol. 2019;100(3):367–8.

26. Mirdita M, Schutze K, Moriwaki Y, Heo L, Ovchinnikov S, Steinegger M. ColabFold: making protein folding accessible to all. Nat Methods. 2022;19(6):679–82.

27. Penzes JJ, Soderlund-Venermo M, Canuti M, Eis-Hubinger AM, Hughes J, Cotmore SF, et al. Reorganizing the family Parvoviridae: a revised taxonomy independent of the canonical approach based on host association. Arch Virol. 2020;165(9):2133–46.

28. Bloom ME, Best SM, Hayes SF, Wells RD, Wolfinbarger JB, McKenna R, et al. Identification of aleutian mink disease parvovirus capsid sequences mediating antibody-dependent enhancement of infection, virus neutralization, and immune complex formation. J Virol. 2001;75(22):11116–27.

29. Alexandersen S. Acute interstitial pneumonia in mink kits: experimental reproduction of the disease. Vet Pathol. 1986;23(5):579–88.

30. Anistoroaei R, Krogh AK, Christensen K. A frameshift mutation in the LYST gene is responsible for the Aleutian color and the associated Chediak-Higashi syndrome in American mink. Anim Genet. 2013;44(2):178–83.

31. Bloom ME, Race RE, Hadlow WJ, Chesebro B. Aleutian disease of mink: the antibody response of sapphire and pastel mink to Aleutian disease virus. J Immunol. 1975;115(4):1034–7.

32. Oie KL, Durrant G, Wolfinbarger JB, Martin D, Costello F, Perryman S, et al. The relationship between capsid protein (VP2) sequence and pathogenicity of Aleutian mink disease parvovirus (ADV): a possible role for raccoons in the transmission of ADV infections. J Virol. 1996;70(2):852–61.

33. Eshak MIY, Rubbenstroth D, Beer M, Pfaff F. Diving deep into fish bornaviruses: Uncovering hidden diversity and transcriptional strategies through comprehensive data mining. Virus Evol. 2023;9(2):vead062.

34. Chen S, Zhou Y, Chen Y, Gu J. fastp: an ultra-fast all-in-one FASTQ preprocessor. Bioinformatics. 2018;34(17):i884–i90.

35. Wood DE, Lu J, Langmead B. Improved metagenomic analysis with Kraken 2. Genome Biol. 2019;20(1):257.

36. Lu J, Rincon N, Wood DE, Breitwieser FP, Pockrandt C, Langmead B, et al. Author Correction: Metagenome analysis using the Kraken software suite. Nat Protoc. 2024.

37. Li H, Handsaker B, Wysoker A, Fennell T, Ruan J, Homer N, et al. The Sequence Alignment/Map format and SAMtools. Bioinformatics. 2009;25(16):2078–9.

38. Prjibelski A, Antipov D, Meleshko D, Lapidus A, Korobeynikov A. Using SPAdes De Novo Assembler. Curr Protoc Bioinformatics. 2020;70(1):e102.

39. Antipov D, Raiko M, Lapidus A, Pevzner PA. Metaviral SPAdes: assembly of viruses from metagenomic data. Bioinformatics. 2020;36(14):4126–9.

40. Hyatt D, Chen GL, Locascio PF, Land ML, Larimer FW, Hauser LJ. Prodigal: prokaryotic gene recognition and translation initiation site identification. BMC Bioinformatics. 2010;11:119.

41. Chen Y, Ye W, Zhang Y, Xu Y. High speed BLASTN: an accelerated MegaBLAST search tool. Nucleic Acids Res. 2015;43(16):7762–8.

42. Sonnhammer EL, Eddy SR, Durbin R. Pfam: a comprehensive database of protein domain families based on seed alignments. Proteins. 1997;28(3):405–20.

43. Gurevich A, Saveliev V, Vyahhi N, Tesler G. QUAST: quality assessment tool for genome assemblies. Bioinformatics. 2013;29(8):1072–5.

44. Quinlan AR, Hall IM. BEDTools: a flexible suite of utilities for comparing genomic features. Bioinformatics. 2010;26(6):841–2.

45. Rozewicki J, Li S, Amada KM, Standley DM, Katoh K. MAFFT-DASH: integrated protein sequence and structural alignment. Nucleic Acids Res. 2019;47(W1):W5–W10.

46. Minh BQ, Schmidt HA, Chernomor O, Schrempf D, Woodhams MD, von Haeseler A, et al. IQ-TREE 2: New Models and Efficient Methods for Phylogenetic Inference in the Genomic Era. Mol Biol Evol. 2020;37(5):1530–4.

47. Blum M, Andreeva A, Florentino LC, Chuguransky SR, Grego T, Hobbs E, et al. InterPro: the protein sequence classification resource in 2025. Nucleic Acids Res. 2025:53(D1):D444–D56.

48. Meng EC, Goddard TD, Pettersen EF, Couch GS, Pearson ZJ, Morris JH, et al. UCSF ChimeraX: Tools for structure building and analysis. Protein Sci. 2023;32(11):e4792.

49. Abramson J, Adler J, Dunger J, Evans R, Green T, Pritzel A, et al. Accurate structure prediction of biomolecular interactions with AlphaFold 3. Nature. 2024;630(8016):493–500.

50. Wakeley PR, Johnson N, McElhinney LM, Marston D, Sawyer J, Fooks AR. Development of a real-time, TaqMan reverse transcription-PCR assay for detection and differentiation of lyssavirus genotypes 1, 5, and 6. J Clin Microbiol. 2005;43(6):2786–92.

