## Supplemental figures for "A novel genetically distinct *Amdoparvovirus* in *Sorex araneus* in the United Kingdom highlights an unexplored ancestral link"

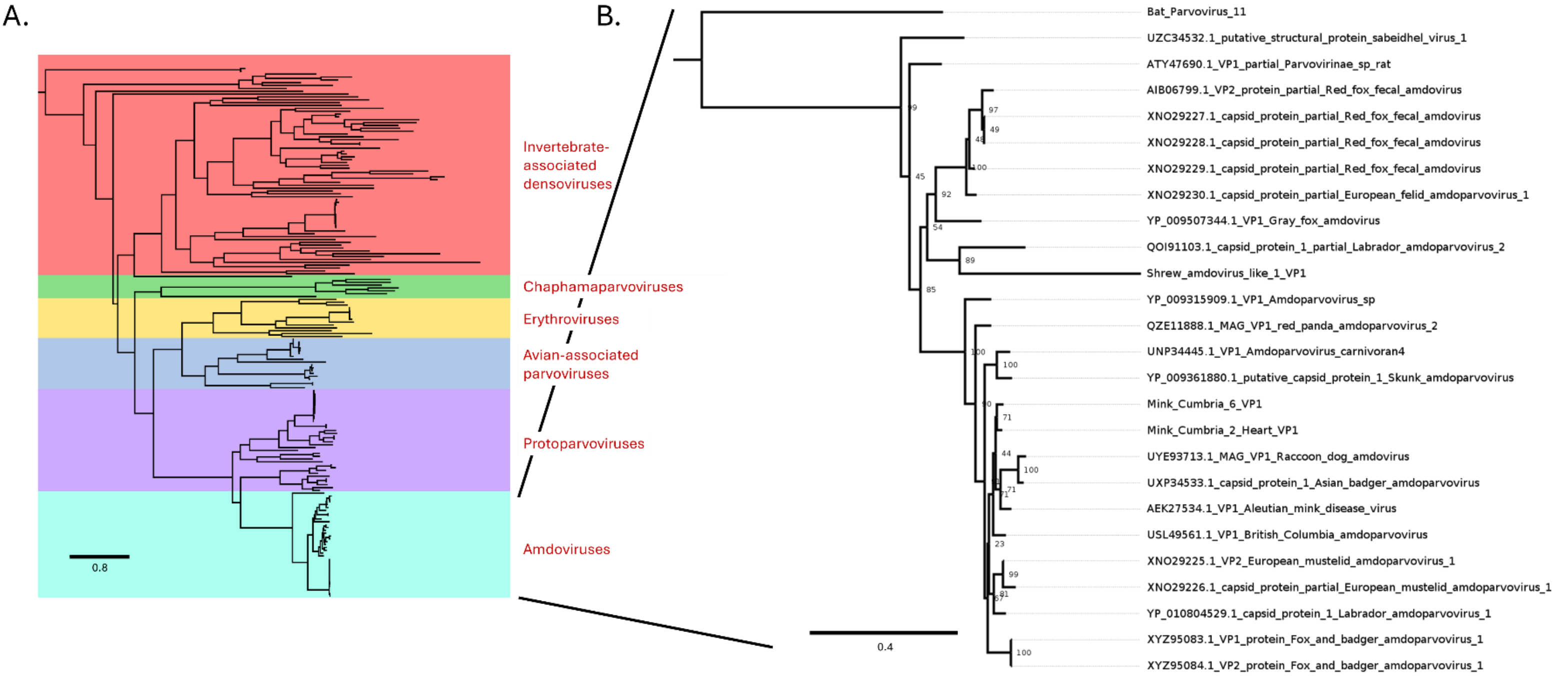


**Supp 1: Phylogenetic tree of VP1 sequences. A. Parvoviridae phylogenetic VP1 tree** showing all parvovirus ref-seq VP1 protein rooted to Bos taurus papillomavirus L1 protein (AZG02908.1). Model used: Q.PFAM+F+R6. **B. Cropped amdovirus clade showing where identified shrew, bat and mink sequences lie** alongside previously reported ref-seq VP1 protein sequences. Phylogenetic tree was rooted with canine parvovirus VP1 (WQF67426.1), 1000 bootstraps, Model used RTREV+I+R3. Putative bat parvovirus 11 was generated through SRA data (SAMN36440297). Last accessed 02/09/2025.

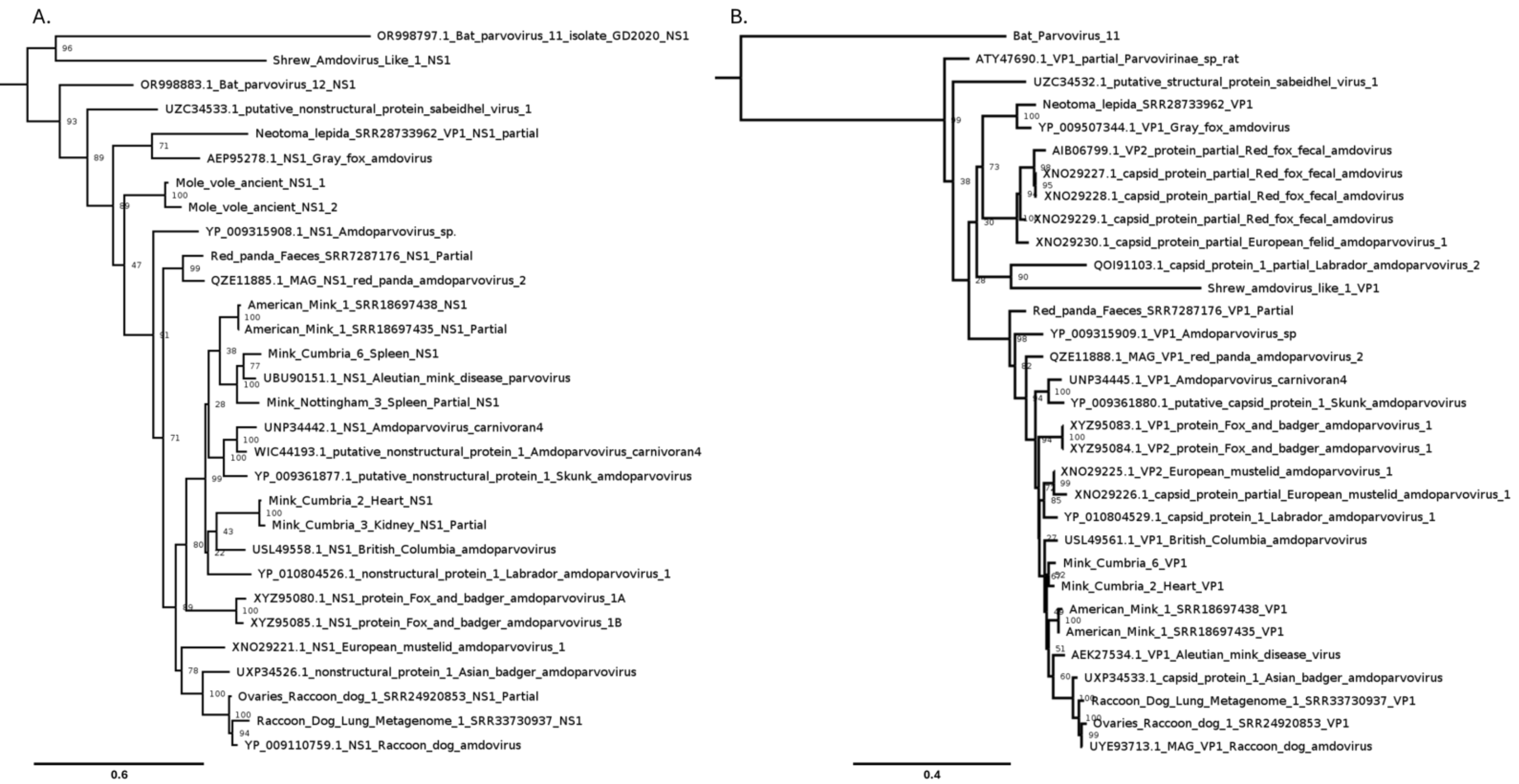


**Supp 2: Phylogenetic trees of NS1 (A) and VP1 (B) sequences including partial SRA mined sequences**. Both rooted to canine parvovirus **(A)** WQF67428.1 **(B)** WQF67426.1 and populated with amdovirus ref-seq and SRA mined sequences. 1000 bootstraps, Models: **(A)** Q.insect+F+G4 **(B)** rtREV+I+R3. Last accessed 01/12/2025.

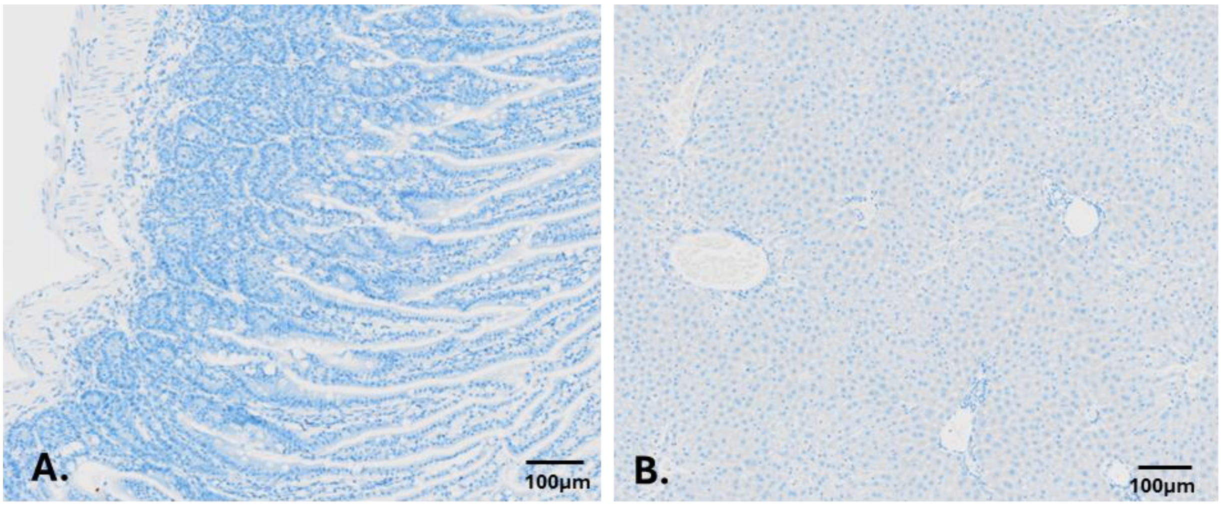


**Supp 3: Negative DapB cISH control sections. A. Intestine. B. Liver** showed no signal. Scale bars are shown. **Abbreviations: cISH;** Chromogenic in situ hybridisation.
